# Nemacounter: A user-friendly software to accurately phenotype SCN cysts

**DOI:** 10.1101/2024.07.07.602381

**Authors:** Joffrey Mejias, Djampa K. L. Kozlowski, Jackson Goshon, Thomas R. Maier, Thomas J. Baum

## Abstract

The soybean-cyst nematode (SCN; *Heterodera glycines*) is one of the most destructive pests affecting soybean crops. Effective management of SCN is imperative for the sustainability of soybean agriculture. A promising approach to achieving this goal is the development and breeding of new resistant soybean varieties. Researchers and breeders typically employ exploratory methods such as Genome-Wide Association Studies or Quantitative Trait Loci mapping to identify genes linked to resistance. These methods depend on extensive phenotypic screening. The primary phenotypic measure for assessing SCN resistance is often the number of cysts that form on a plant’s root system. Manual counting hundreds of cysts on a given root system is not only laborious but also subject to variability due to individual assessor differences. Additionally, while measuring cyst size could provide valuable insights due to its correlation with cyst development, this aspect is frequently overlooked because it demands even more hands-on work. To address these challenges, we have created Nemacounter, an intuitive software designed to detect, count, and measure the size of cysts autonomously. Nemacounter boasts a user-friendly graphical interface, simplifying the process for users to obtain reliable results. It enhances productivity by delivering annotated images and compiling data into csv files for easy analysis and reporting.

## Introduction

Soybeans rank among the most widely cultivated crops, with Brazil and the United States being the largest producers. In the 2021-2022 growing season, soybean production reached 399.50 million metric tons serving as a significant source of both human food and animal feed (http://www.worldagriculturalproduction.com/crops/soybean.aspx). The ability of legumes like soybeans to form symbiotic relationships with rhizobia bacteria diminishes the need for external fertilizer application making soybean very useful in crop rotations^1,2^. One of the major threats to soybean cultivation is the soybean cyst nematode (SCN; *Heterodera glycines*), a highly destructive phytopathogen^3^. There are only relatively few resistance sources deployed into available soybean cultivars and their effectiveness is slowly eroded by the development of novel virulence phenotypes in SCN populations. Consequently, there is an urgent need to breed additional resistant germplasms and to develop novel pest management strategies^4^. Such novel management avenues will require extensive SCN research, including high volume phenotyping.

Cyst nematodes are filiform worms (juvenile stage 2; J2) that invade the roots of soybean plants and reach the vascular cylinder, adopting a sedentary lifestyle. They induce the formation of a feeding structure by manipulating root cells to alter their cell walls and fuse iteratively with adjacent cells, forming a syncytium^5^. Nematodes feed exclusively on this structure after its formation, withdrawing nutrients and energy necessary for its further development. These nematodes reproduce sexually, with fertilized females producing hundreds of eggs. The egg-laden female hardens into a cyst on the surface of infected roots. These cysts easily dislodge into the soil, and new juvenile nematodes hatch from the eggs within^5,6^. Therefore, the quantity and size of cysts found on a plant’s roots are crucial measures of infection success by SCN.

New sources of resistance to cyst nematodes can be discovered through both classical and reverse genetic screening methods. Forward screening techniques, such as Genome-Wide Association Studies (GWAS) and Quantitative Trait Loci (QTL) analysis, are potent tools for identifying genes that confer resistance to pathogens^7–9^. The effectiveness of GWAS and QTL analysis is heavily dependent on the precision of infection measurements^10^. This requirement for accurate infection assessment is equally important in reverse genetic screening in which specific genes are evaluated for their ability to influence infection levels. Such assessments are critical for determining whether a particular gene plays a role in the interaction between SCN and soybean plants.

At present, most scientists assess cyst numbers manually under a binocular microscope. Manual counting of cysts is not only labor intensive and time-consuming but also very prone to inconsistencies, underlying the need for more efficient and uniform methods for assessing infection rates. Accurate phenotyping of SCN infection success is vital in breeding efforts and also to evaluate the effects of host genetic mutations or environmental conditions. Subtle variations in cyst numbers might be overlooked if the phenotyping method lacks precision and accuracy. Additionally, despite its importance, measuring each cyst’s size to detect developmental delays and reproductive success is rarely done in nematology research due to the laborious and time-intensive nature of the task if performed manually, except in cases where the difference is starkly apparent^11^.

To address these challenges, we have developed “Nemacounter” a user-friendly software utilizing the pre-trained neural network *You Only Look Once* ^12,13^,*14 version 5 (YOLOv5) combined with the Segment Anything Model* (SAM). Our software precisely and accurately identifies, counts, and measures the size of each SCN cyst in a 1040x1040 pixel size image taken on a white background without the need for specialized equipment. Nemacounter is straightforward to use and generates output that includes labeled images and an Excel spreadsheet containing detailed information about the count and size of each cyst per image as well as the average pixel cyst size per picture, along with SD and SE. The accuracy of cyst detection using the Nemacounter software has been evaluated at ∼96%, which is roughly the same rate achieved by a trained human. Moreover, to improve upon this 4% error rate, Nemacounter allows the user to optionally review automatically annotated pictures one-by-one, and easily adjust the annotation manually before the segmentation. The software can be installed and run locally on a Windows or Linux computer using a standard central processing unit (CPU), offering flexibility for users, for which we provide a detailed installation procedure. The Nemacounter software can be downloaded here: https://github.com/DjampaKozlowski/NemaCounter and we provide an installation manual and utilization manual as supplementary data.

## Results

### Nemacounter: a high-precision SCN cyst counting and sizing software

We introduce Nemacounter, a user-friendly software designed using a custom Python script (Fig. 1). Nemacounter leverages the YOLOv5-xl Object Detection pre-trained neural network for object (cyst) detection and the SAM^15^ for precise size extraction of each detected cyst (Fig. 1). We choose this set-up after comparing different approaches. We first explored the possibility of using various versions of the YOLO algorithm for object detection, particularly focusing on recent iterations such as YOLOv5, YOLOv8 and YOLO-NAS. The YOLOv8 and YOLO-NAS advanced versions demonstrate enhanced efficiency compared to YOLOv5-xl, as highlighted in previous research (https://github.com/ultralytics/ultralytics). However, they exhibit reduced stability during training and are unsuitable for processing images of 1040x1040 pixels on an A100 GPU due to their substantial computational requirements. Attempts to mitigate this issue by resizing images to dimensions such as 640x640 using those recent versions resulted in a noticeable decline in detection performance when comparing YOLOv8-m using 640x640 image input to YOLOv5-xl using 1040x1040 image input (Fig. S1). Consequently, the latest versions of YOLO were deemed unsuitable for our purposes.

**Figure 1.**
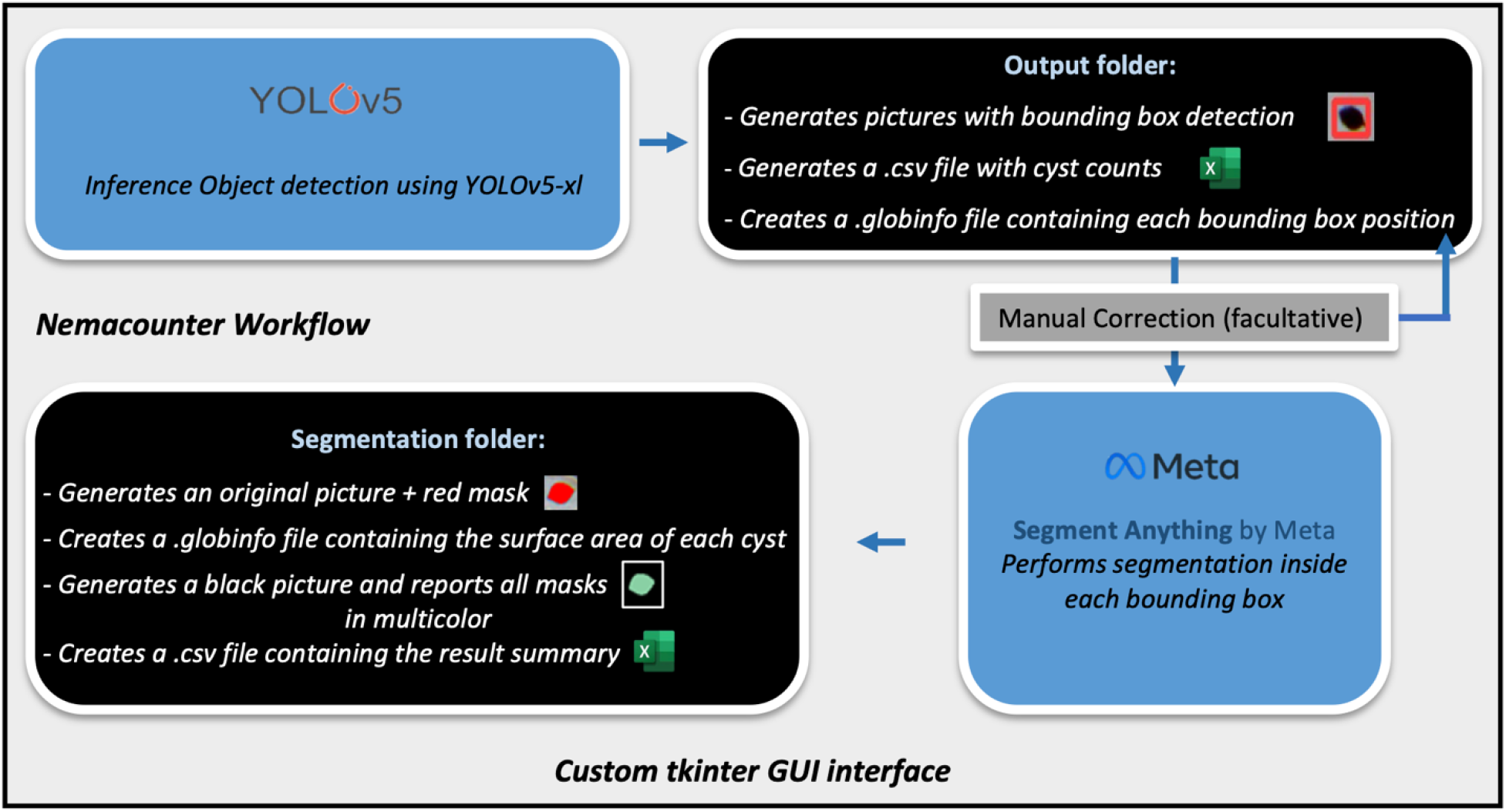
Nemacounter workflow. Nemacounter is based on a custom python script using customtkinder library for the GUI interface. The inference is performed through the YOLOv5-xl Object Detection model and the custom script to create an « output » folder containing the original pictures with the detected bounding boxes in red; a .csv file containing the picture name and the corresponding cyst count as well as .globinfo files containing each bounding box position for each cyst per picture. A manual correction step can be applied using the OpenCV-tool library that modified the previously generated .globinfo file after manual correction. The segmentation is performed through the SAM developed by Meta inside each updated bounding boxes coordinates contained in the glob info file. After segmentation, an output folder called « segmentation » is created and contains: the original pictures with the overlapping mask (in red), the same pictures with random multicolored masks on a black background, and tables that summarizes the results for each picture including (i) the cyst count, (ii) size for each cyst per picture, and (iii) the average pixel cyst size as well as each corresponding SD and SE per picture.

Our objective was also to develop a software capable of accurately quantifying the size of each cyst. Initially, we considered employing the YOLOv5-xl Instance Segmentation variant, which utilizes polygonal annotations to delineate objects, thereby directly extracting masks corresponding to cyst areas. We annotated cysts using polygons and trained this model using 640x640 resized pictures (Fig. S2). However, this approach yielded suboptimal results in cyst detection compared to the YOLOv5-xl Object Detection model (Fig. 2) but has the advantage of providing size measurements for each cyst. To address these limitations, we used a hybrid method which involves applying SAM within the bounding boxes identified by the YOLOv5-xl Object Detection model (Fig. 3). This new approach retains the high-quality detection rate of the YOLOv5 Object Detection model and also allows cyst size extraction by SAM. Upon comparing the two methodologies, it was evident that this combined approach produced masks of superior resolution compared to the standalone YOLOv5 Instance Segmentation method (Fig. 2), and therefore was implemented in the Nemacounter software.

**Figure 2.**
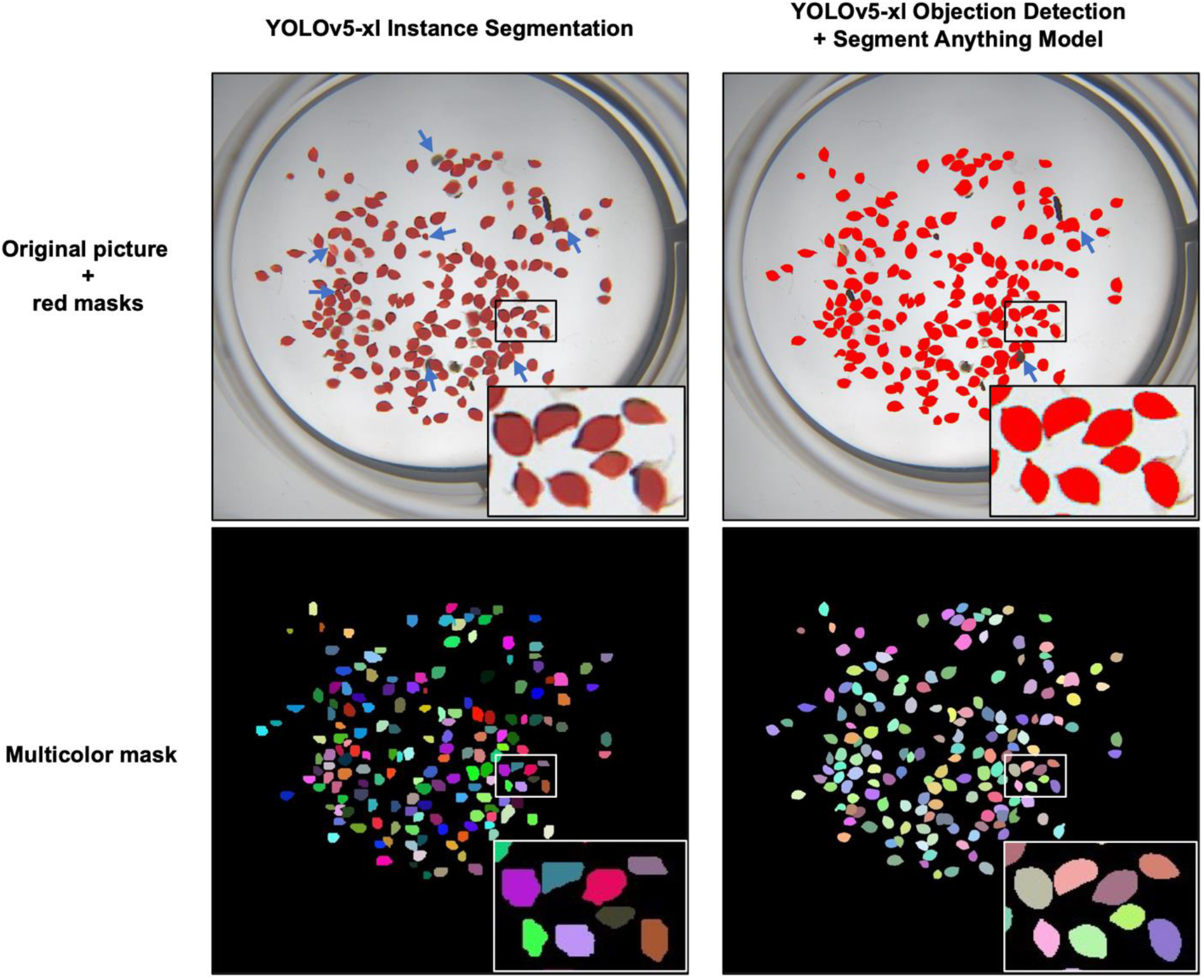
Comparison between the standalone YOLOv5-xl Instance Segmentation model and the combination of YOLOv5-xl Object Detection + SAM. The figure shows the results after segmentation for both methods. Blue arrows indicate errors made by the YOLOv5-xl Instance Segmentation model, or errors made by the combination of YOLOv5-xl Object Detection model + SAM models.

**Figure 3.**
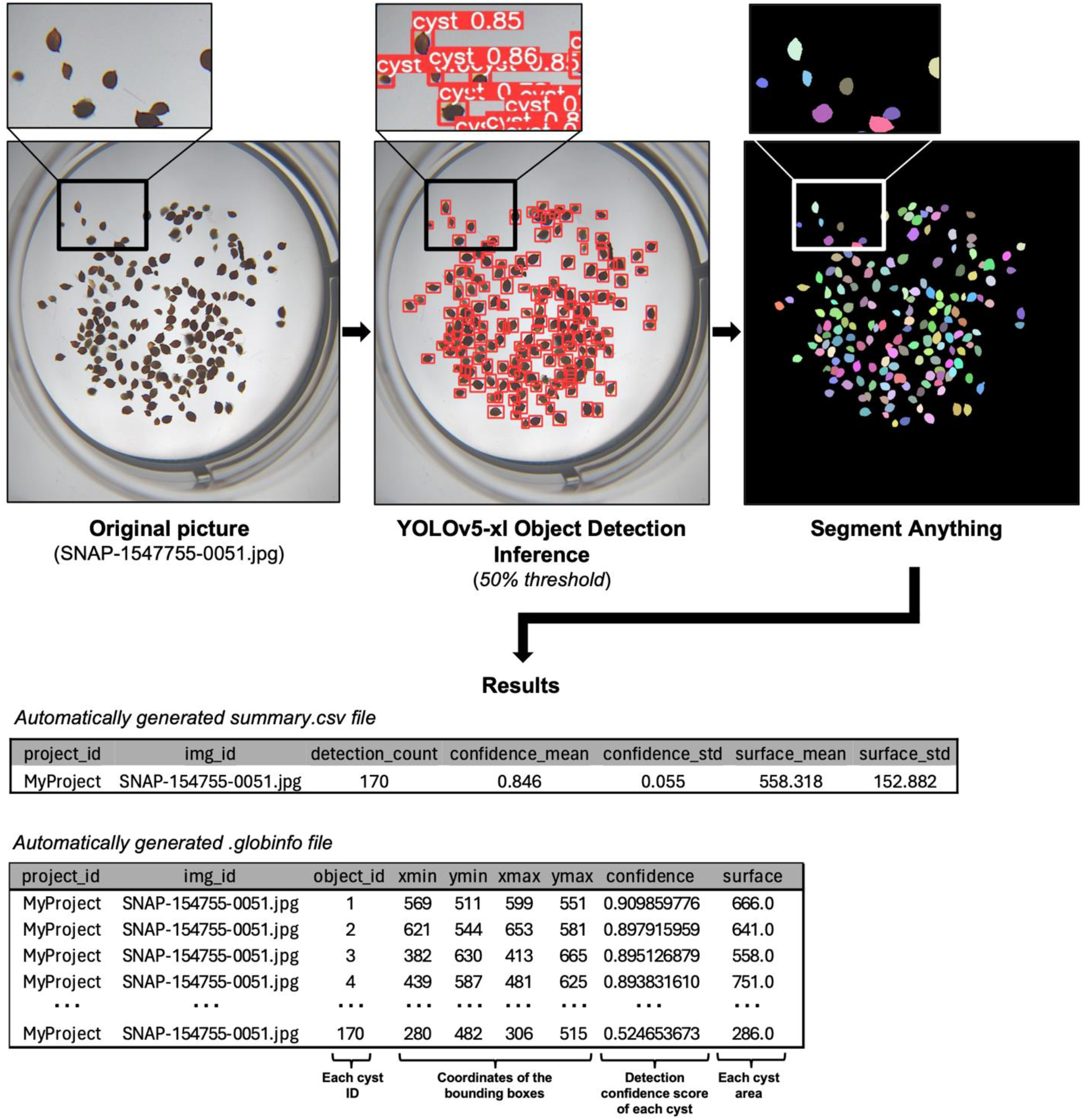
Example of output results obtained after using the Nemacounter software on a picture never seen before. On the left, the original picture before inference and a zoomed area. In the middle, picture containing the bounding boxes that were detected by inference using the YOLOv5-xl Object Detection model. The zoomed area contains the confidence score of each cyst. The original picture is shown without labels and confidence scores. On the right, the random multicolored masks produced by the script on a black background after segmentation. At the bottom, an example of the automatically generated tables for this picture (SNAP-1547755-0051).

### Neural network training and performance metrics

Utilizing a NVIDIA A100 GPU, the YOLOv5-xl Object Detection neural network was trained on a dataset composed of 183 pictures (162 for training and 21 for validation) for a total of 12,005 cysts manually annotated (Fig. S3; Fig. S4). At the 386^th^ epoch, considered the best weight, the model displayed a box loss of 0.046908, indicating minimal error in bounding box location predictions (Fig. 4). The object loss stood at 0.4917, reflecting the model’s accuracy in object detection. Notably, the precision metric reached 0.98128, illustrating high accuracy in positive instance prediction, and a recall of 0.96167, highlighting the model’s ability to identify relevant instances (Fig. 4). The Mean Average Precision (mAP) at an Intersection over Union (IoU) threshold of 0.5 was an impressive 0.98726, showcasing accuracy in object detection with moderate overlap. However, at varying IoU thresholds (0.5 to 0.95), the mAP dipped to 0.71971 indicating a performance decrease when cysts overlapped too much (Fig. 4).

**Figure 4.**
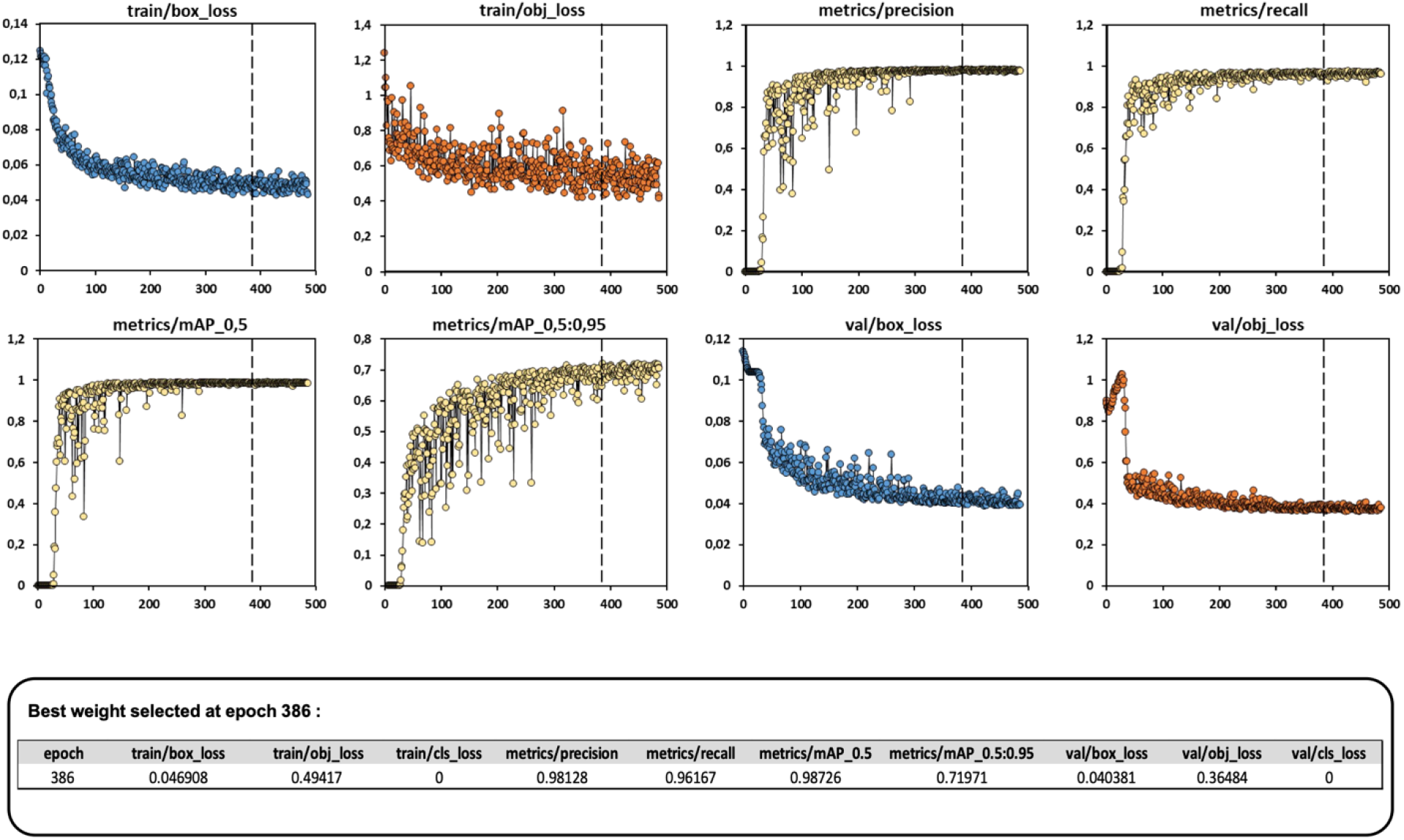
Training results of the YOLOv5-xl Object Detection pre-trained neural network. Each parameter resulting from the training such metrics, precision or recall are plotted. The dotted line represents the best epoch 386. Below are the final parameters kept for the best weight at 386 epoch. Precision is the ratio of true positive detections to all detections made, while recall is the ratio of true positive detections to all actual objects present. The curves illustrate the trade-off between precision and recall at various confidence thresholds. The mean average precision (mAP) at an Intersection over Union (IoU) threshold of 0.5 (mAP@0.5) and the mean average precision averaged over IoU thresholds from 0.5 to 0.95 (mAP@0.5:0.95) are shown, which measure the model’s overall detection performance across different classes and IoU thresholds.

### User interface and functionality

Nemacounter’s GUI, built on customtkinter package, allows users to select input folders containing cyst images in various formats (JPG, JPEG, PNG, TIFF). The software automatically processes results to a chosen output folder. Users can adjust detection thresholds via a slider, with the default setting at 0.5, as well as the IoU threshold with the default setting at 0.3 to filter objects identified as cysts with over 50% probability and 0.3 IoU threshold. The default input image size is 1040x1040 pixels. Confidence levels on each cyst as well as the mean confidence are scored in the generated .globinfo file (Fig. 3).

During inference, the processing time per image averages 300 ms. After inference, the output folder includes images with detected cysts, and corresponding bounding box coordinates stored in .globinfo file. Users can review results by going into this output folder while the software is open and adjust results by re-running inferences with different thresholds, creating additional output folders as needed.

### Manual correction, segmentation process, and results summary

If users identify incorrectly annotated cysts, they can manually correct annotations using Nemacounter’s OpenCV-based tool, which allows the user to draw or remove bounding boxes on each picture (Fig. S5). Following the optional manual corrections, segmentation is initiated to extract cyst sizes within each updated bounding box’s coordinates present in the new .globinfo file generated by clicking on the “Segmentation” button. This process generates a “Segmentation” folder containing visual representations of segmented cysts with red masks on original images, along with a randomly multicolored mask image to verify individual cyst segmentation on a black background (Fig. 3). Additionally, a .globinfo file and a summary.csv files are generated, listing the number, size, and statistical analysis (average, standard deviation, standard error) of cysts in each image (Fig. 3).

### Validation of Nemacounter’s object detection performance

To rigorously evaluate Nemacounter’s detection capabilities without applying any manual corrections afterwards, we conducted tests using eight images previously unseen by the neural network. This set comprised four ‘clean’ cyst samples and four ‘dirty’ cyst samples that also contained root debris (Fig. 5), each manually annotated. The Nemacounter script was employed for detection and segmentation on both datasets. Based on the coordinate of each mask extracted automatically for these pictures, we calculate the centroids of each cyst using the following methodology:

**Figure 5.**
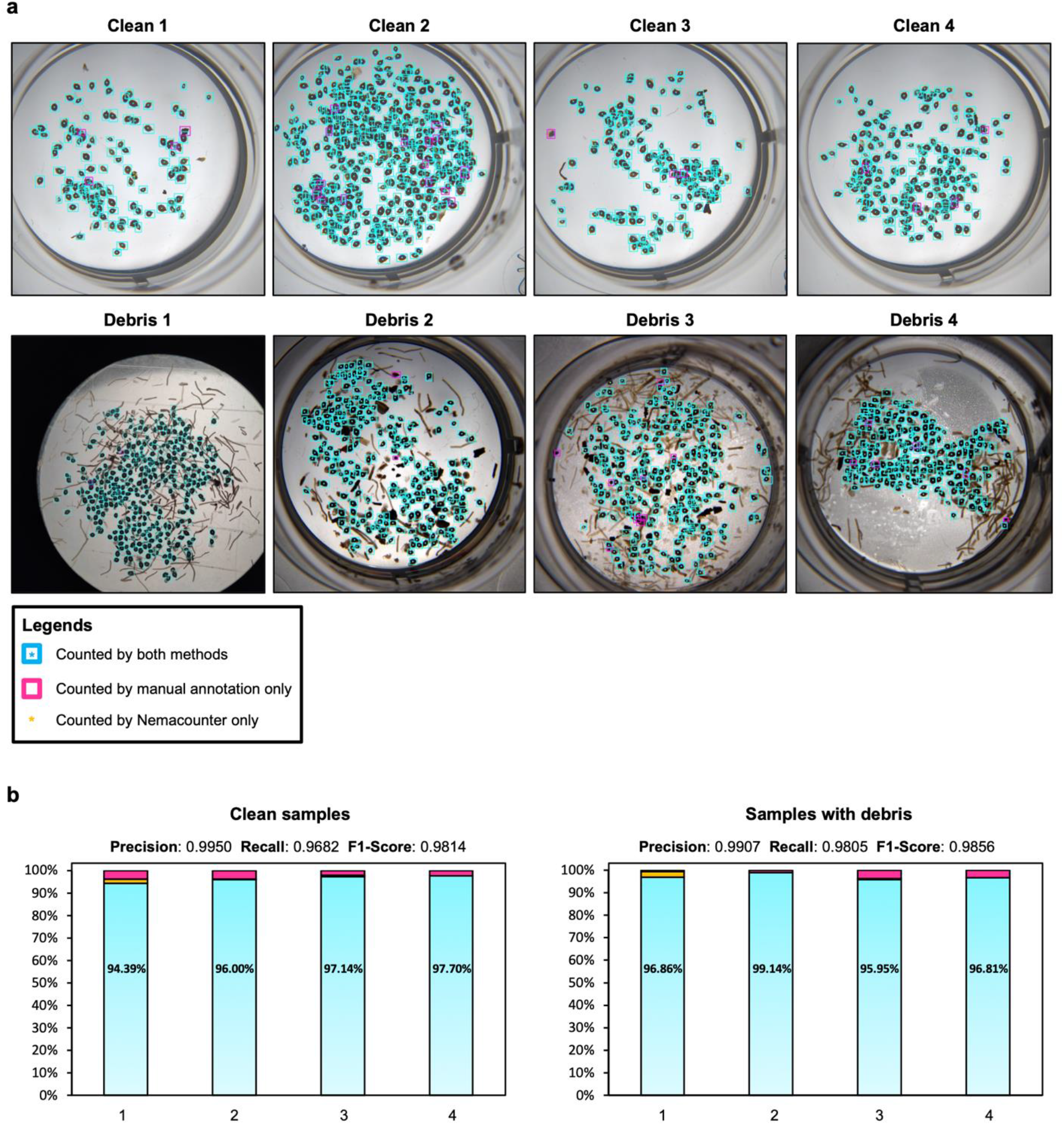
Evaluation of the Nemacounter’s ability to correctly detect cysts on unseen pictures. (**a**) Four different pictures containing cyst without debris (upper row) and four pictures containing cyst with debris (lower row). Visual comparison of manual annotation versus automatic annotation using Nemacounter. Below the legend shows the signs used to designate the cysts that were counted by both methods (cyan asterisks and box), by only Nemacounter (yellow asterisks), or by manual annotation alone (magenta box). (**b**) Histogram showing for each picture the percentage of cyst counted by both methods (cyan) as well as the corresponding percentage, by Nemacounter (yellow), or by manual annotation (magenta). Counts for the three categories were pooled for the four clean pictures or pictures containing debris to calculate precision score, recall score and F1-score, which are shown for each pool above the corresponding histogram.

#### Centroids calculation

Utilizing OpenCV’s cv2.moments function, we calculated the moments of each cyst contour that correspond to the weighted averages of the pixel intensities in the contour of the cyst. A contour represents the continuous line forming the boundary of the cyst in the image. The zeroth spatial moment, denoted as M[‘m00’], corresponds to the area of the contour. The first order spatial moments, M[‘m10’] and M[‘m01’], are integral in determining the centroid’s coordinates. The centroid’s X-coordinate (*C*_*x*_) is computed as the ratio of the first order moment along the x-axis to the zeroth moment, i.e., *C*_*x*_ *= M[‘m10’]/ M[‘m00’]*. Similarly, the Y-coordinate (*C*_*y*_) is obtained by dividing the first order moment along the y-axis by the zeroth moment, i.e., *C*_*y*_ *= M[‘m01’]/ M[‘m00’]*. The calculated coordinates, (cx, cy), represent the centroid’s position on the image (Fig. S6).

#### Comparison of centroid detected automatically to manual annotation

We utilized a custom script to determine whether the centroids of automatically identified cysts fell within the manually annotated bounding boxes. If a given centroid detected automatically fell into a bounding box manually annotated, it means that the software is in accordance with the manual annotation for a given cyst in a picture. Each centroid was aligned to the nearest bounding center and uniquely assigned, preventing multiple assignments to the same box. This method enabled a direct visual comparison between cysts counted by Nemacounter with those identified manually, or those identified only by the Nemacounter (Fig. S6). Distinct categories were recognized: cysts counted by both methods, those counted exclusively by manual annotation, and cysts detected solely by Nemacounter (Fig. 5a). The comparative analysis revealed that Nemacounter aligns with manual annotations with an average accuracy of 95% across both clean and dirty samples, which is similar to a human error rate when counting hundreds of cysts on a given picture (Fig. 5b). We pooled the count of the four pictures corresponding to the clean sample and calculated the Precision score (0.9950), the Recall score (0.9682), and the F1-Score (0.9814). We performed a similar analysis with the four pictures corresponding to the sample with debris and observed a Precision score of 0.9907, a Recall score of 0.9805, and F1-Score of 0.9856. On its own, the Nemacounter performed with 96% accuracy, which can be further improved as needed by supplementing automatic annotations with manual correction inputs, thereby filling the 4% discrepancy if necessary (Fig. S5).

### Analysis of cyst size measurement accuracy

To determine Nemacounter’s efficacy in discerning cysts of different sizes, we used ImageJ to manually sort cysts by size into three categories under a light microscope (small, medium, and large) and placed them into a 12 well-plate to take pictures (Fig. 6a). We generated three images for each category, allowing for a thorough examination of Nemacounter’s ability to accurately capture and reflect these size differences using the SAM technology. This approach aimed to validate whether the pixel size extraction feature of SAM could effectively detect and categorize varying cyst sizes in the samples, which will be particularly useful to assess if a given condition was affecting cyst development. We first ran these nine pictures (three pictures per category) through the Nemacounter and directly plotted the results from the automatically generated .globinfo file and observed a clear distribution of the three size categories, demonstrating Nemacounter’s accuracy to extract cyst size (Fig. 6a-b). Subsequently, we compared the cyst sizes that were manually determined using ImageJ software to the mask extracted by Nemacounter. We manually measured each cyst present in each size group for a total of 323 cysts using ImageJ or Nemacounter (Fig. 6a,c). We observed a clear correlation between the size of each cyst for each manually or automatically labeled picture (Fig. 6d-f). Although, we observed a small discrepancy between the two methods where the ImageJ method show higher cyst area compared to Nemacounter, this discrepancy merely appears to be due to the way the contours of cyst have been drawn in ImageJ, which slightly overestimates the actual area, whereas Nemacounter tends to not completely extract cyst head and tails (Fig. 6c). We then grouped all cysts extracted by both methods and performed a linear regression which showed a R^2^= 0.9978, demonstrating a high correlation between manual size measurements and the Nemacounter (Fig. 6g). Overall, these results demonstrate that Nemacounter precisely extracts the size of each cyst in a repeatable manner essential for comparing two conditions in an experimental set-up designed to assess SCN infectivity and development.

**Figure 6.**
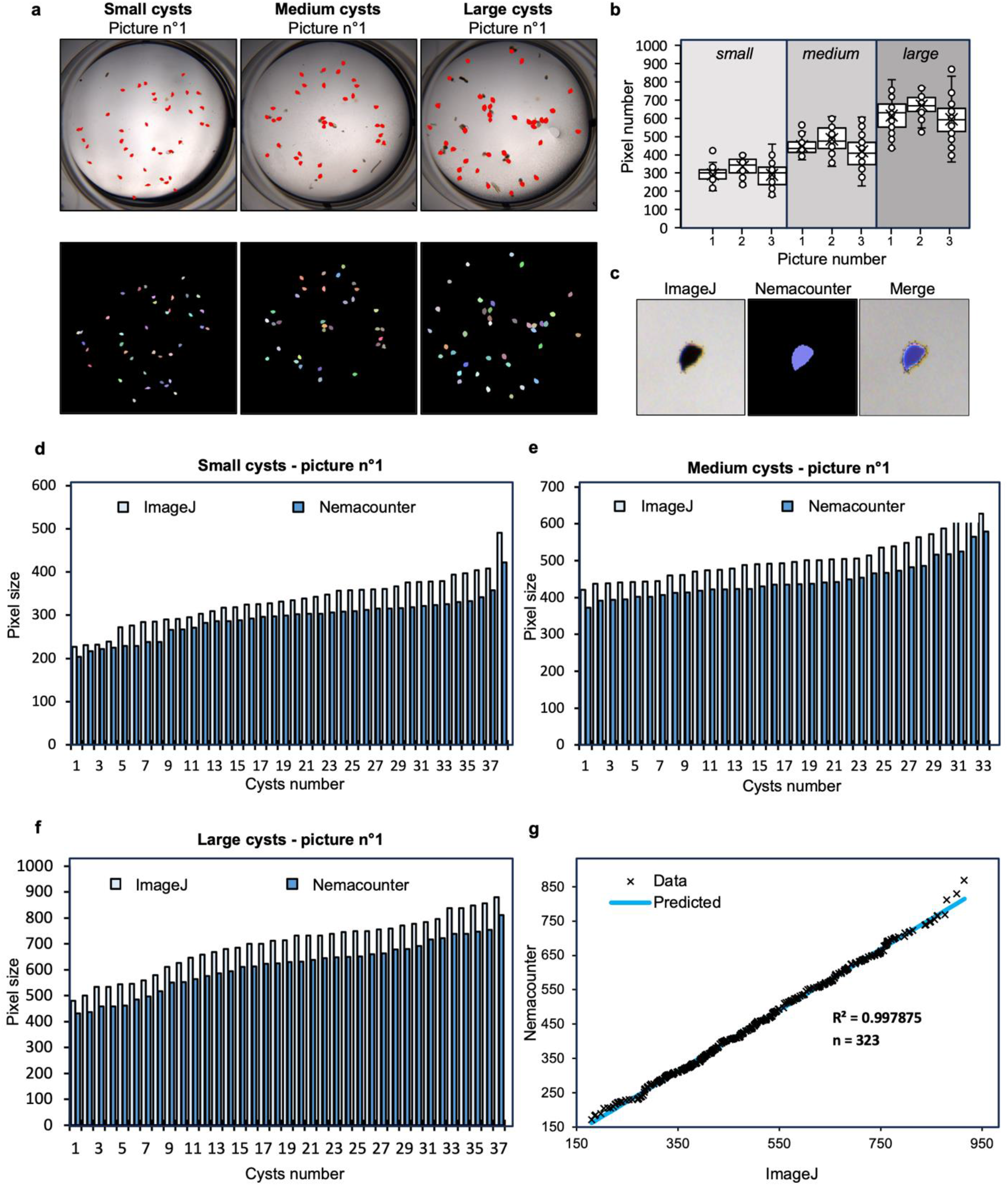
Evaluation of Nemacounter’s ability to extract cyst size. (**a**) Cysts were physically sorted by size into three categories: small, medium and large. One representative original picture for each category is displayed with a red mask (upper row) or the random multicolored mask displayed on black background (lower row). (**b**) Whisker-plot of each cyst’s size measured by Nemacounter in each picture are shown. Three pictures per size category were analyzed. (**c**) Representative illustration of a cyst size measurement performed by hand using ImageJ (left) or automatically extracted using Nemacounter (middle), and the merge of the two methods (right). (**d-f**) Histograms showing cyst size in each picture measured using ImageJ or the Nemacounter. For each category, one picture is displayed (**d** = small cysts; **e**= medium cysts and **f** = large cysts). (**g**) Linear regression of all the cyst size (pixel size) from the nine pictures showing the correlation between ImageJ and Nemacounter cyst size measurement.

## Discussion

In this study, we developed Nemacounter to help nematologists to accurately count and extract the size of cysts in infection experiments without any labor-intensive work or required skills in bioinformatics. Nemacounter is an easy-to-use software, offering a user-friendly interface that processes multiple images simultaneously and in various formats. We initially compared the latest YOLO models (YOLOv5, YOLOv8 and YOLO-NAS). Unfortunately, the YOLOv8 and YOLO-NAS neural networks are too big and therefore cannot be trained using a 1040x1040 pixel size input picture on a A100 GPU. The need for resizing the picture to a smaller scale (640x640 on a A100 GPU) renders the detection less efficient than using the YOLOv5-xl Object Detection Model which reaches high performance (mAP50 = 0.98) and therefore does not require improvement or the use of a larger neural network. We combined the detection of the YOLOv5-xl Object Detection model and SAM to accurately count and subsequently extract cyst size with precision. We compared this method with the YOLOv5-xl Instance Segmentation model and found that our method outperforms the instance segmentation model when both are trained using the best parameters on a A100 GPU (Fig. 2). This poorer quality results observed using the YOLOv5-xl Instance Segmentation is certainly due to the need of resizing the pictures to 640x640. Upon comparison, the masks resolution extracted have much better resolution using YOLOv5-xl Object Detection + SAM (Fig. 2). Indeed, the SAM model can segment any object thanks to a training performed using over 1 billion masks from 11 million pictures, making it extremely precise to segment any area.

We developed a custom method to compare the efficiency of detection of the Nemacounter versus manual annotation (Fig. 5a; Fig. S6). The output metrics obtained after training gives us a good idea of the performance of the model but only vaguely informs us on the nature of false positives and false negatives created during inference. Therefore, to visually assess the nature of false positive and false negative errors that are created by the Nemacounter, we calculated the centroids of each cyst from the mask generated by the software and compared them to manually annotated bounding boxes. If a centroid falls into a bounding box, it is assigned to it and therefore cannot be assigned to another box. If two centroids or more fall into a bounding box, the centroid that is closest to the center of the bounding box is assigned to it. This process allows us to generate pictures and assess visually the nature of the error as well as the rates of false positives and false negatives produced by the Nemacounter (Fig. 5b; Fig. S6). In our case, an average of 4% error rate total is observed, which is in accordance with the recall score (0.96) and is mostly due to overlapping cysts (Fig. 5). Although these errors can be manually corrected using the manual correction option in Nemacounter, we advise users to shake plates containing the cysts gently before taking pictures to minimize such overlapping cases and obtain the best automated performance.

We next assessed if the Nemacounter could extract cyst size in an accurate and repeatable manner. To do so, we have manually sorted cysts into three size categories and plotted the results directly from the output excel table generated by the Nemacounter. We observed a clear repartition of the three groups, confirming the robustness of the approach (Fig. 6b). To go further, we used ImageJ to manually extract the size of each cyst and compared these values with the results provided by the Nemacounter (Fig. 6c). We observed that the manual annotation (using polygons) overestimates the actual cyst area whereas the Nemacounter tends to not fully extract the area corresponding to the head and the tail of the cyst (Fig. 6c). Despite this discrepancy, we observed a correlation of 99.78% between the two methods demonstrating that the Nemacounter can be used to accurately measure cyst size (Fig. 6g).

In the current iteration of Nemacounter, we have integrated a YOLOv5-xl Object Detection model followed by SAM, which allows us to integrate an intermediate step of manual correction. After initial inference, any necessary adjustments to bounding box coordinates can be easily made by overwriting the .globinfo file, streamlining the process before proceeding to segmentation with SAM. This flexibility in correction is a significant advantage over a standalone instance segmentation model such as the YOLO-xl Instance Segmentation in which it would have been much more difficult to implement this option. Moreover, Nemacounter creates an excel file summarizing the whole analysis as well as the pictures containing the masks (red masks on original picture, or masks in a random multicolor pattern on black background) (Fig. 3). The GUI developed on customtkinter provided a simple interface where the user chooses the input folder containing the pictures to analyze and select an empty folder on his computer where the result will be saved.

Previously, two studies have employed neural networks to detect SCN cysts. The first study developed a custom neural network using MATLAB and WEKA with specified parameters such as texture to recognize which species of cyst was present, including *Globodera pallida, Globodera rostochiensis*, and *Heterodera schachtii*^*16*^, however this approach does not provide cyst number or area for users. Another study used the U-Net, ResNet50 and ResNet101 instance segmentation models for the specific purpose of detecting SCN cysts in soil samples^17^. The set-up requires the use of a high-quality camera and applies a stitching method on multiple tiles from split pictures before processing to be able to detect cysts on a large paper containing soil sample^17^. Nemacounter does not require a high-resolution camera. Indeed, any camera able to take pictures at 1040x1040 pixel size can be used with our software. Furthermore, we have developed a GUI interface, added a manual correction step, all on a user-friendly interface. Other neural networks have been used to recognize different genera of nematodes such as EfficientNetV2M that allows recognition of 11 genera of nematodes^18^ or the DenseNet121, which was used for the development of NemaNet, that recognizes larvae stages of *Helicotylenchus dihystera, Heterodera* *glycines* (J2), *Meloidogyne incognita* (J2), *Pratylenchus* *brachyurus, Rotylenchulus* *reniformis*^*19*^. To enhance the utility of nematology research tools, nematologist could extend the capabilities of existing software to include the detection and classification of various stages of phytoparasitic nematode development, such as eggs, and juvenile stages J2 through J4, as well as males. Advanced models like YOLOv8 or YOLO-NAS could be used to categorize these numerous stages with high precision. However, such sophisticated operations would necessitate the use of multiple, powerful GPUs concurrently for the training phase. Researchers have also developed an approach to track sugar-beet cyst nematode (BCN, *Heterodera schachtii*) development when inoculated *in vitro* for high-throughput screening of infected *Arabidopsis thaliana* plants^20^. This study provided a low-cost and open-source method for nematode phenotyping that included nematode size as a scorable parameter and a method to account for phenotypic variation of the host. This methodology was based on threshold-mediated object isolation and contrast coloration to detect female nematodes and also provided a 3D printable model to set-up the system in the lab^20^. Implementation of neural networks could further improve that method in the future.

The application of YOLO models to video analysis presents exciting opportunities for future development, particularly in tracking J2 nematodes during migration assays. Typically, these assays involve placing J2s in a square box filled with pluronic gel where chemo-attractants are placed at opposite ends^21,22^. While scientists traditionally count the number of nematodes at each end post-experiment to compare chemoattractant effectiveness, a YOLO-based video tracking system could revolutionize this process. Such a system would allow for continuous tracking and counting of J2s as they reach either pole, offering a dynamic timeline and more nuanced insights into migration patterns.

Now that the Nemacounter script is publicly available, it can be adapted to help analysis of a wide array of objects beyond nematology. Users can utilize the script’s functionality by simply updating the path to the best weight file coming from another training. The GUI and output features, such as inference, manual correction, segmentation, and excel table creation, will therefore remain applicable to any other objects. Indeed, annotating pictures or training a model using YOLOv5-xl on Google Colab does not require any coding skills. This flexibility paves the way for diverse applications for biologists with no bioinformatic skills such as bacterial colony-forming unit counting, fungal spore counting or fungal lesion disease quantification on leaves, and more. In summary, Nemacounter allows accurate counting of cysts as well as cyst size extraction, and reports results. Nemacounter will help nematologists to analyze their infection assays involving SCN more rapidly and in a repeatable manner.

## Materials and Methods

### Collection and photography of soybean cyst nematode samples

Soybeans were grown in a controlled chamber at 24°C at 45-65% relative humidity and a 16h/8h light/dark cycle. After about 2-3 weeks, soybean plants were transferred into 8-inch containers filled with a mixture of two-parts sand to one-part field soil that had previously been steam sterilized. Two days later, each plant was infected with 1000 J2 SCN TN10. Thirty days after infection, the SCN cysts were collected in a manner similar to that described previously^23^. Finally, purified cysts were placed either on glass Petri dishes or into 12 well-plates. To obtain “clean samples” or “samples containing root debris”, we used successive sucrose gradient cleanups. Pictures were taken using a Zeiss Stemi SV11 microscope with AxioCam HRc and Axiovision SE64 V4.9.1 software to snap pictures of 1040x1040 resolution on white background. To obtain pictures with varying contrasts, we additionally took pictures using the upper LEDs to light the cyst from above.

### Dataset compilation and artificial augmentation

Our dataset encompassed a diverse range of high-contrast images falling into three groups for comprehensive analysis: 1) SCN cysts in glass Petri dishes, 2) SCN cysts in 12-well plates, 3) SCN cysts displayed in varied colorations in 12-well plates (Fig. S3). This assortment of images enhanced the model’s ability to recognize cysts under varying conditions. The training set included 54 images, as well as a validation set comprising 21 images, totaling 12,005 manually annotated cysts (Fig. S4). The complete dataset is accessible on the Roboflow website at: https://universe.roboflow.com/iowa-state-university-cwvqa/cystnewboundingboxv2. To artificially augment the dataset, we applied various picture transformations to the original training dataset: vertical/horizontal flipping, 90° rotation, 20% cropping, 25% gray-scaling, and ±25% brightness adjustment, effectively tripling the number of training images to 162 (Fig. S4).

### Training using YOLOv8-m Object Detection model

We utilized the YOLOv8-m Object Detection neural network, the medium sized YOLOv8 model for object detection, comprising 218 layers and approximately 2.58 million parameters. Training was conducted on a NVIDIA A100 40 GB RAM GPU using Google Colab’s Roboflow notebook. Training parameters were set as follows:

!yolo task=detect mode=train model=yolov8m.pt data={dataset.location}/data.yaml epochs=3000 imgsz=640 plots=True project=‘/content/drive/My Drive/bestmodel’ name=‘YoloV8run’

We used a 640x640 resizing parameter for training and 16 batches as parameters, which are the maximum parameters that can be used on this model using an A100 GPU (Fig. S1a). The best epoch was selected at 153 to prevent overfitting the dataset as no significant improvements were noted in the subsequent 50 epochs. Inference was run using an IoU equal to 0.3 and an 640 x 640 input image size.

### Training using YOLOv5-xl Instance Segmentation model

We trained the YOLOv5-xl Instance Segmentation neural network using polygon annotation available on the Roboflow website at: https://universe.roboflow.com/iowa-state-university-cwvqa/cyst-detectors-area. YOLOv5-xl is the largest model for instance segmentation of YOLOv5, comprising 330 layers and approximately 8.8 million parameters. Training was conducted on a NVIDIA A100 40 GB RAM graphical processing unit (GPU) using Google Colab’s Roboflow notebook. Training parameters were set as follows:

python train.py --img 640 --batch 20 --epochs 3000 --data [dataset.location]/data.yaml –cfg ./models/custom_yolov5x.yaml --weights ‘‘ --project “/content/drive/MyDrive/bestmodel” --name yolov5xseg_results –cache

We used a 640x640 resizing parameter for training and 20 batch as parameters, which are the maximum parameters that can be used on this model using an A100 GPU (Fig. S2). The best epoch was selected at 741 to prevent overfitting the dataset as no significant improvements were noted in the subsequent 100 epochs. Inference was run using an IoU equal to 0.3 and an 640 x 640 input image size.

### Training using YOLOv5-xl Object Detection model

We utilized the YOLOv5-xl Object Detection neural network, the largest YOLOv5 model for object detection, comprising 233 layers and approximately 7.25 million parameters. Training was conducted on a NVIDIA A100 40 GB RAM GPU using Google Colab’s Roboflow notebook (Fig. 4). Training parameters were set as follows:

python train.py --img 1040 --batch 80 --epochs 3000 --data [dataset.location]/data.yaml –cfg ./models/custom_yolov5x.yaml --weights ‘‘ --project “/content/drive/MyDrive/bestmodel” --name yolov5xobj_results --cache

The decision to use 1040x1040 pixel images without resizing was intentional to preserve intricate details for the neural network’s learning process. Training halted at epoch 386 to prevent overfitting, as no significant improvements were noted in the subsequent 100 epochs. The resulting “cystmodel.pt” is available at: https://iastate.box.com/s/e9kfpkjkrfpye0wgje205023xpcmcgvs.

### Inference using YOLOv5-xl Object Detection model, manual correction, and segmentation process

Nemacounter integrates a custom Python script, offering an intuitive tkinter-based user interface. It harnesses the YOLOv5-xl Object Detection model and the possibility to manually correct the annotation if needed, followed by segmentation using the SAM developed by Meta. Inference starts with the YOLOv5-xl Object Detection model, using the following command:

python “C:\Users\Username\yolov5\detect.py” --weights “C:\Users\Username\yolov5\cystmodel.pt” --source “[input_folder]” --project “[output_folder]” -- name output --iou-thres 0.3 --conf-thres [conf_thres] --img-size [img_size] [hide_labels_and_conf_arg] --save-txt

For inference, an IoU threshold of 0.3 was chosen to balance detecting closely situated cysts while avoiding redundant counts. The input image size was maintained at 1040x1040 pixels, consistent with the training dataset. Users are recommended to use a similar white background and pixel size for optimal performance with Nemacounter. The GUI allows the user to modify the arguments in this command without modifying the script.

The manual correction can be applied if needed on each picture using a user-friendly interface where the user can draw new bounding boxes (holding left-click to draw) or remove bounding boxes (by pressing the “R” key when the mouse is hovering on a given bounding box). The modified manual correction is saved by pressing the “S” key which will close the current picture and open the next one. The manual correction window can therefore be quit by pressing the “Esc” key.After inference (and optionally after manual correction), the segmentation is performed inside each bounding box by clicking in “Run Segmentation”. The segmentation uses the SAM biggest model weight. The “sam_vit_h” is available at: “https://dl.fbaipublicfiles.com/segment_anything/sam_vit_h_4b8939.pth“ or “https://iastate.box.com/s/akpql0jlvbd5mmw26e2lgya4ul9h1xua“.

## Acknowledgments

We would like to thank Melissa Bredow for critical review of the manuscript.

## Author Contributions

JM conceived of the project. JM, JG and TRM conducted the experiments. JM and DK wrote the software. TJB provided guidance and financial support. JM wrote the manuscript with feedback from all authors.

## Data Availability Statement

All training datasets are available on Roboflow website at : https://universe.roboflow.com/iowa-state-university-cwvqa/cystnewboundingboxv2 and https://universe.roboflow.com/iowa-state-university-cwvqa/cyst-detectors-area. Nemacounter is available on github at: https://github.com/DjampaKozlowski/NemaCounter. The SAM model is available at: https://dl.fbaipublicfiles.com/segment_anything/sam_vit_h_4b8939.pth or https://iastate.box.com/s/akpql0jlvbd5mmw26e2lgya4ul9h1xua. The YOLOv5-xl Object Detection Model is available at: https://iastate.box.com/s/e9kfpkjkrfpye0wgje205023xpcmcgvs.

## Additional Information

This publication contains work funded by the Iowa Agriculture and Home Economics Experiment Station, Ames, IA, supported by Hatch Act and State of Iowa funds and grants from the North Central Soybean Research Program (C00075686/C000838723). The authors declare no competing interests.

## Supplemental Files

**File S1**. How to install Nemacounter.

**File S2**. How to use Nemacounter.

