## Supplemental data for "Nemacounter: A user-friendly software to accurately phenotype SCN cysts"

**a**

**Training of the « YOLOv8-m Object Detection model »**

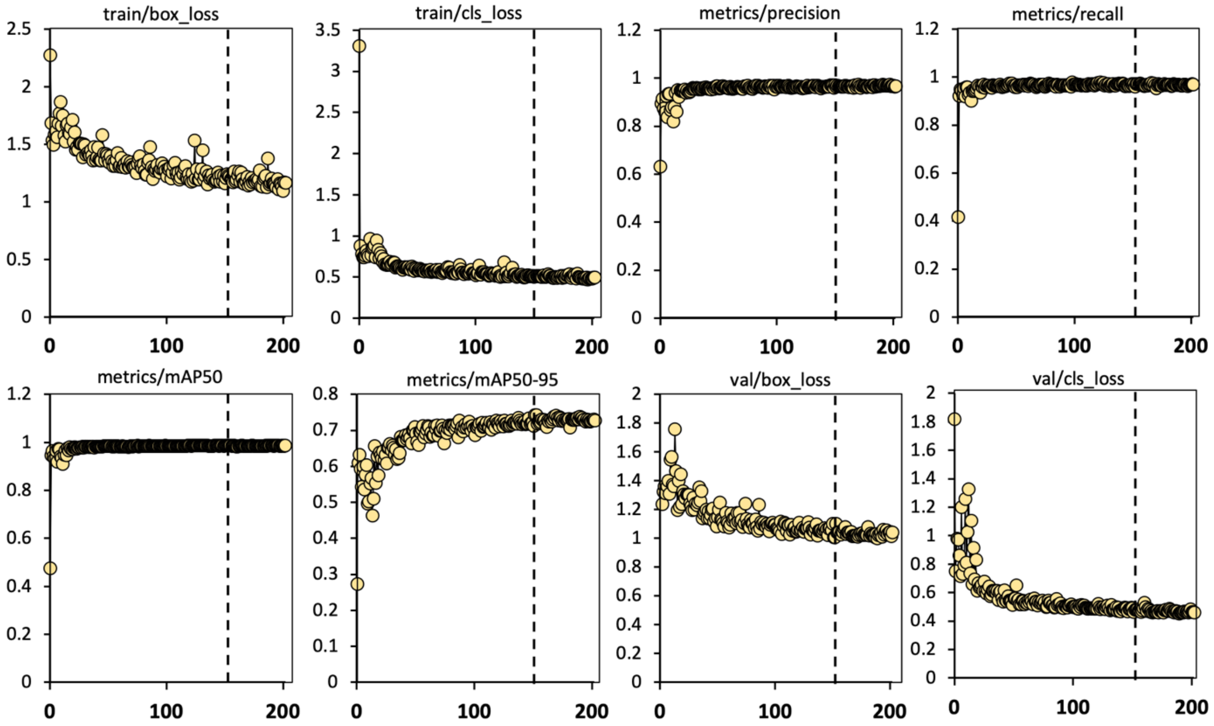


Best weight at epoch 153:

| train/box_loss | train/cls_loss | metrics/precision | metrics/recall | metrics/mAP50 | metrics/mAP50-95 | val/box_loss | val/cls_loss |
|---|---|---|---|---|---|---|---|
| 1,2278 | 0,50992 | 0,96848 | 0,97125 | 0,9887 | 0,71917 | 1,1043 | 0,48614 |

**b**

**YOLOv5xl inference**
Conf=0.5 ; imgsize=1040

**YOLOv8-m inference**
Conf=0.5 ; imgsize=640

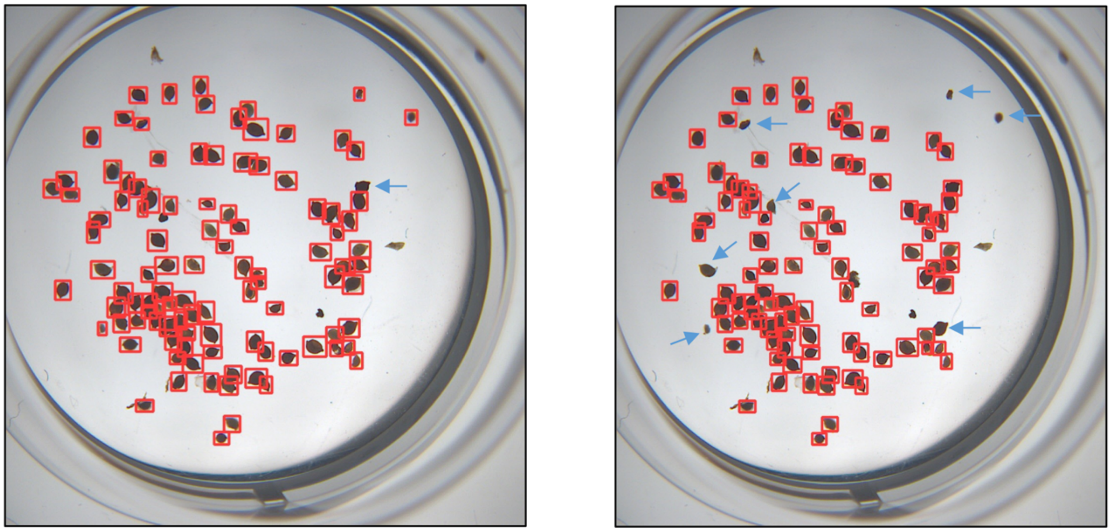

**Supplemental Figure 1**. (**a**) Training results of the YOLOv8-m Object Detection pre-trained neural network on the training dataset. Each parameter resulting from the training such as metrics, precision, or recall are plotted. The dotted line represents the best epoch (153). Below is the final parameters kept for the best weight at 153 epoch. (**b**) Representative comparison between the YOLOv5-xl and the YOLOv8-m Object Detection models. The blue arrows indicate obvious errors in cyst detection.

**Training of the « YOLOv5-xl Instance Segmentation model »**

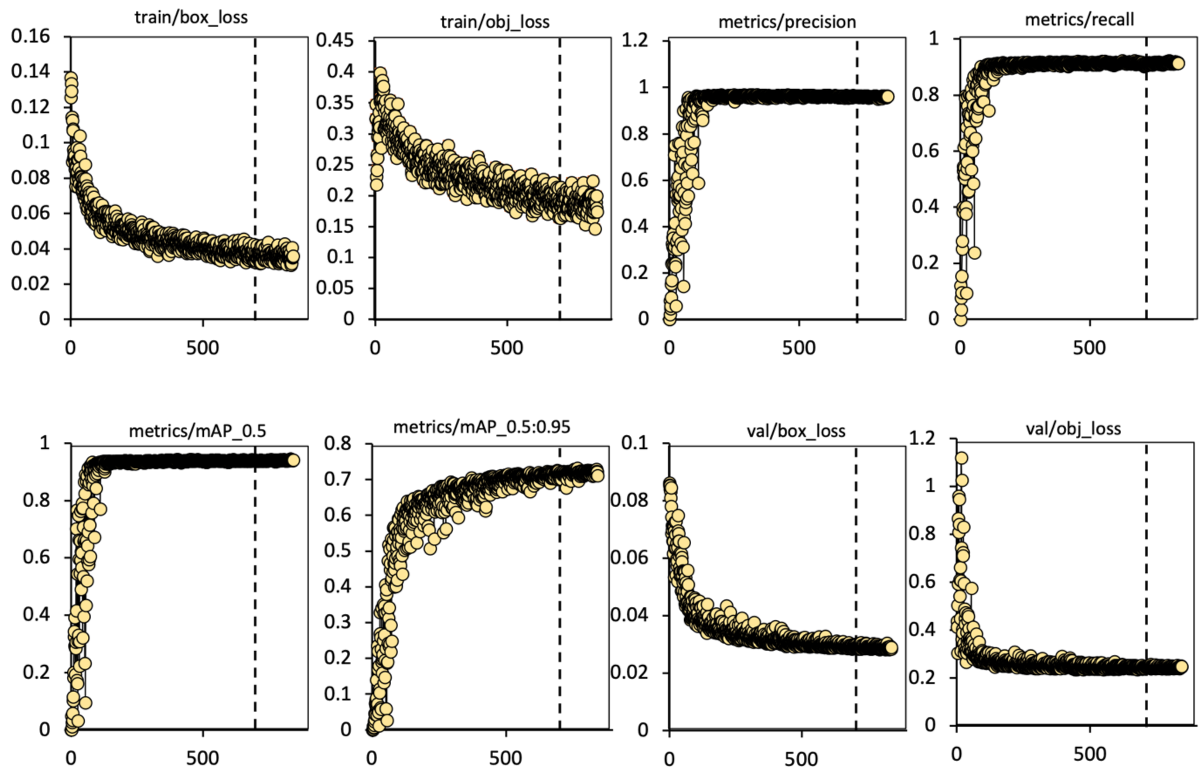


Best weight at epoch 741:

| train/box_loss | train/obj_loss | metrics/precision | metrics/recall | metrics/mAP_0.5 | metrics/mAP_0.5:0.95 | val/box_loss | val/obj_loss |
|---|---|---|---|---|---|---|---|
| 0.036568 | 0.19348 | 0.96251 | 0.90976 | 0.93834 | 0.71978 | 0.028673 | 0.23976 |

**Supplemental Figure 2**. Training results of the YOLOv-xl Instance Segmentation pre-trained neural network on the training dataset. Each parameter resulting from the training such metrics, precision or recall are plotted. The dotted line represent the best epoch (741). Below is the final parameters kept for the best weight at 741 epoch.

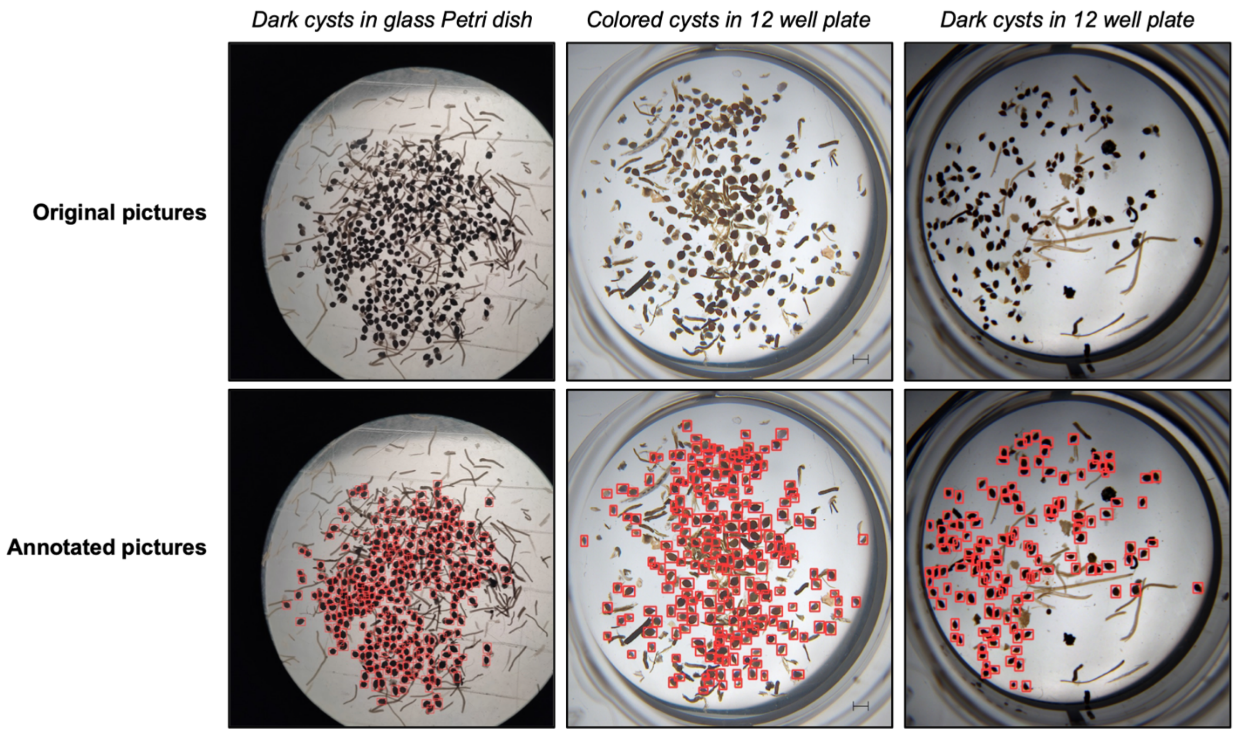


**Supplemental Figure 3**. Dataset used to train the YOLOv5-xl Object Detection model. The dataset contained three picture set-ups: (i) dark cyst in glass Petri dish, (ii) colored cysts in 12 well-plate, and (iii) dark cyst in 12 well-plate. For each set-up, 80% of the pictures were used for training and 20% for validation.

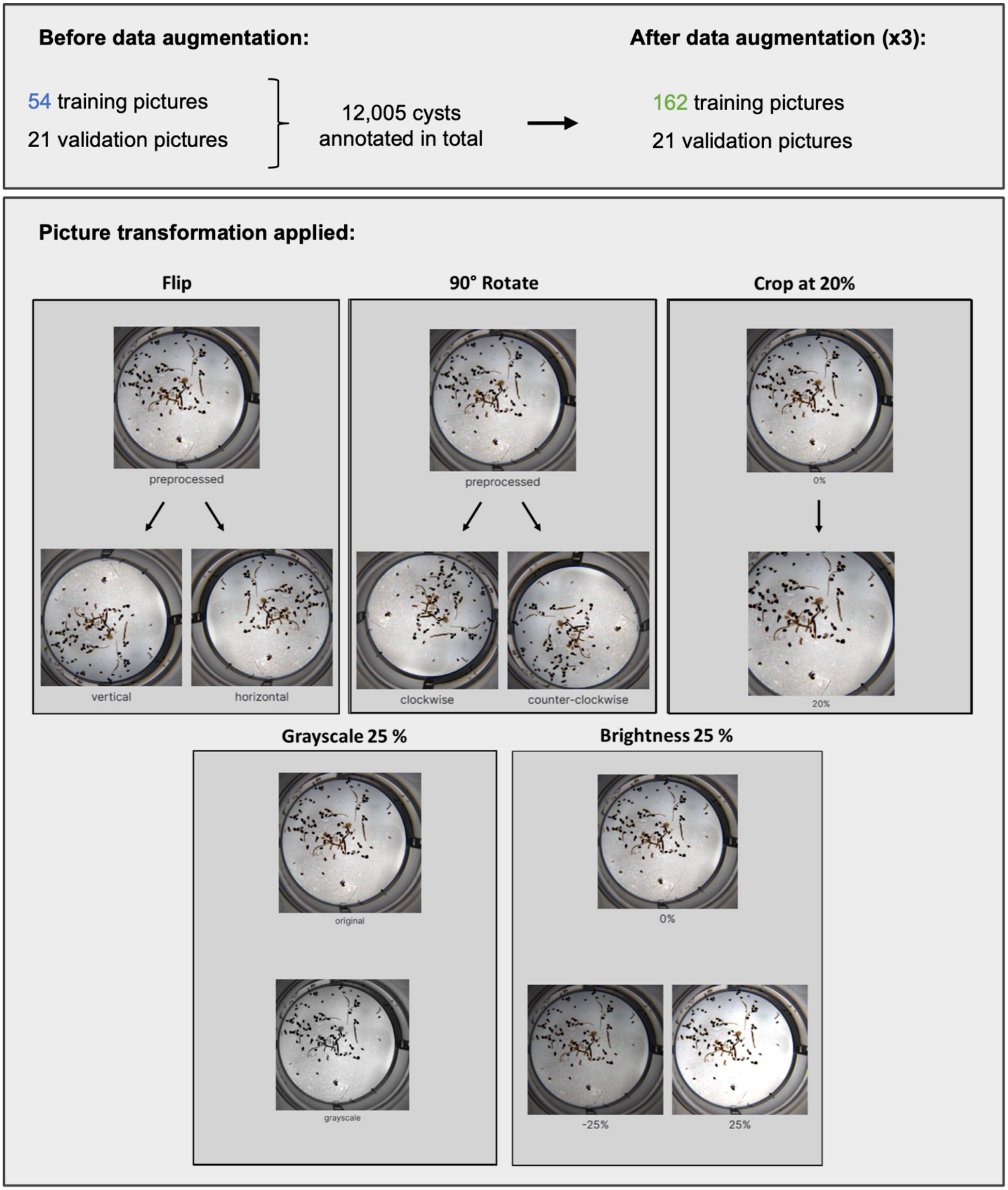


**Supplemental Figure 4**. Artificial augmentation of the training dataset. Figure shows the number of pictures before and after augmentation, as well as the type of transformation, that was applied to the dataset, which includes: flip (vertical/horizontal), 90° rotation (clockwise/counter-clockwise), 20% crop, -/+ 25% grayscale, and -/+ 25% brightness.

**Nemacounter without manual correction**

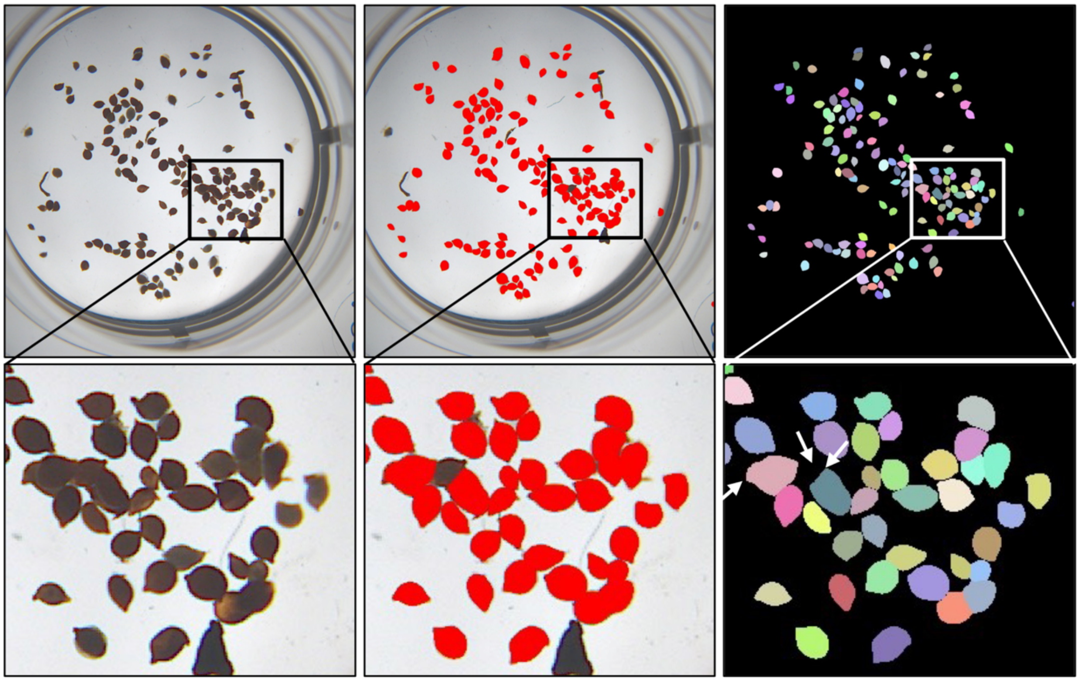

**Nemacounter with manual correction (some bounding boxes are redrawn manually before segmentation)**

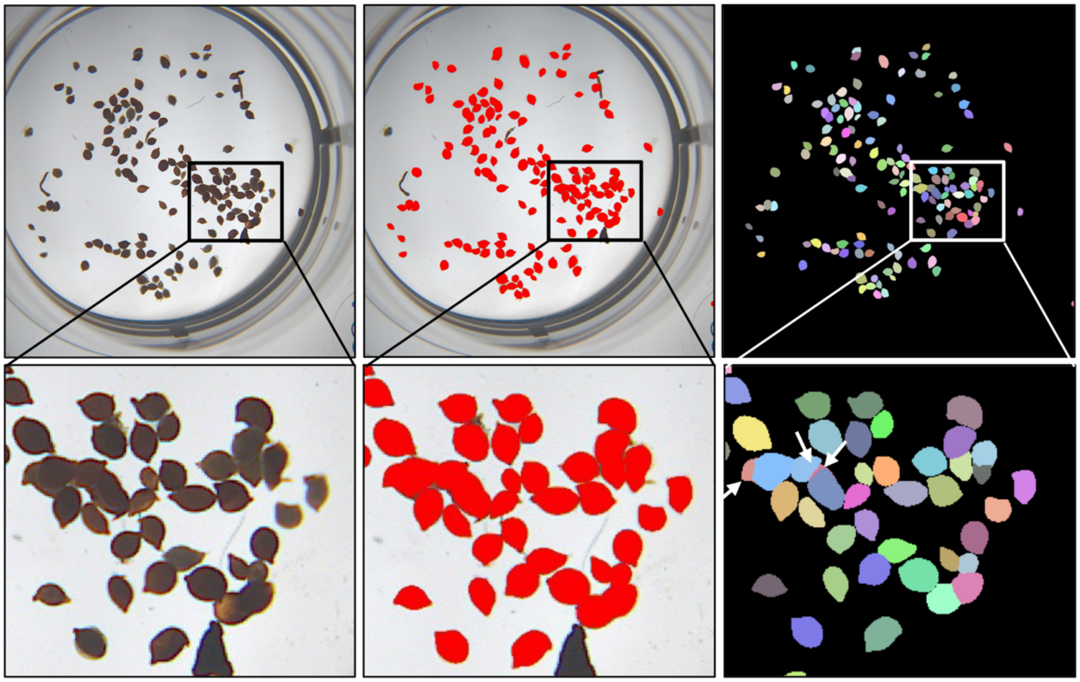

**Supplemental Figure 5**. Example of the manual correction option. From left to right for both top and bottom panels: original picture, original picture with the red mask after segmentation, and the random multicolored masks on a black background. An example of a picture after processing by the Nemacounter without applying manual correction (top row). The same picture after applying manual correction (lower row). The white arrows show the position of missing cysts before and after manual correction.

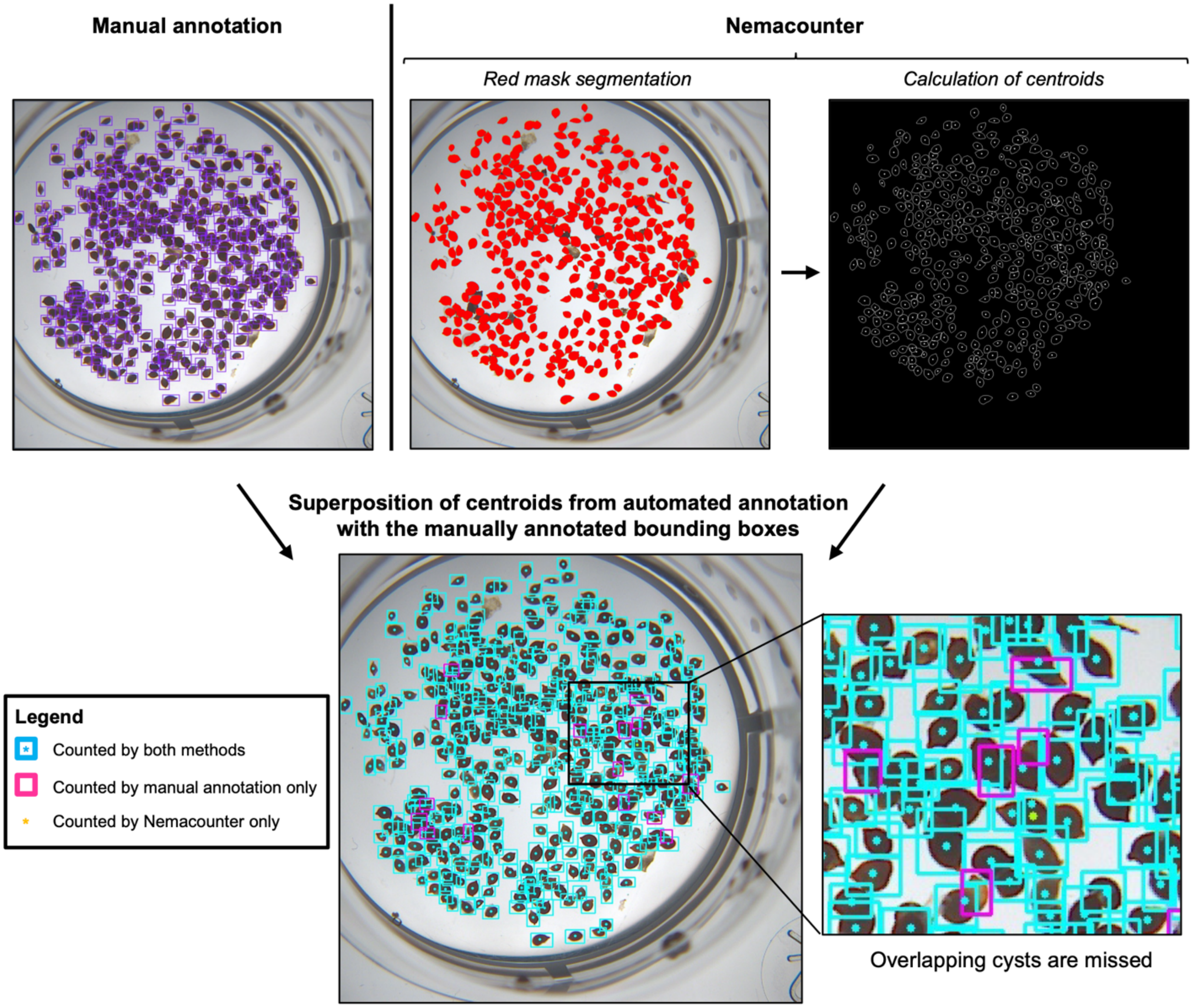


**Supplemental Figure 6**. Example of the custom validation approach for cyst detection. One picture is shown as an example. Shown are the manually annotated picture (top-left), the picture after segmentation by Nemacounter (top-middle), and the picture after centroid calculation. Below, the picture after centroid assignment to each manually annotated bounding box is shown.
