## Supplemental file 1 for "Nemacounter: A user-friendly software to accurately phenotype SCN cysts"

### Installation Instructions of Nemacounter

1. **Download and Install Python for Windows:**

    - If you don't have Python already installed, download and install it from Python's official website --> Window installer (x64 or x32).
    - During installation, do not forget to "Add python to path".

2. **Download and Install Anaconda for Windows:**

    - If you don't have Anaconda already installed, download and install it from Anaconda's official website. Follow the installation instructions and let the anaconda folder be installed in the “Username” folder as per default.

3. **Clone the Nemacounter folder from Github:**
    - Go to https://github.com/DjampaKozlowski/NemaCounter to clone/download the Nemacounter folder on your Desktop.
    - Download the “sam_vit_h_4b8939.pth” model from : https://iastate.box.com/s/akpql0jlvbd5mmw26e2lgya4ul9h1xua and the “cystmodel.pt” from : https://iastate.box.com/s/e9kfpkjkrfpye0wgje205023xpcmcgvs
    - Drap and drop the two downloaded models into the “models” folder located into the Nemacounter folder cloned/downloaded from Github.

4. **Create and Activate the Conda Environment:**

    - Open “Anaconda Prompt” (by searching "anaconda" in your Windows search bar and click on it).
    - Once the terminal is open, create a new environment called "Nemacounter" by copy/pasting the following command:

```
conda create -n Nemacounter python=3.10
```

Press the **“Y”** key follow by “enter” when prompted in the Anaconda window.

- Activate the environment by copy/pasting the following command:

```
conda activate Nemacounter
```

- Change directory to the Nemacounter folder in Anaconda command window by copy/pasting the following command:

```
cd C:\Users\username\Desktop\Nemacounter
```

Make sure to replace `username` by your computer username.

- Install all the required dependencies by copy/pasting the following command:

```
pip install -r requirements_windows.txt
```

or “requirements_linux.txt” if using Linux.

- Wait for all dependencies to be downloaded and installed in the Anaconda prompt command window.
- The Anaconda prompt command window terminal can be close.

#### Run the Application

Now that Nemacounter is installed through Anaconda follow the steps below to run the software each time you need to open it:

1. Open “Anaconda Prompt” (by searching "anaconda" in your Windows search bar and click on it).
2. Activate the environment by copy/pasting the following command:

```
conda activate Nemacounter
```

3. Change directory to the Nemacounter folder in Anaconda command window by copy/pasting the following command:

```
cd C:\Users\username\Desktop\Nemacounter
```

4. Run the software in Anaconda command window by copy/pasting the following command:

```
python Nemacounter_gui.py
```
