## Supplemental file 2 for "Nemacounter: A user-friendly software to accurately phenotype SCN cysts"

### Nemacounter User Guide

#### Overview

Nemacounter is a versatile tool designed for object detection, manual edition, and object segmentation using sequentially YOLOv5 Object Detection, OpenCV library and Segment Anything Model. This guide provides step-by-step instructions on using the software through its three main tabs: Object Detection (A), Manual Edition (B), and Object Segmentation (C).

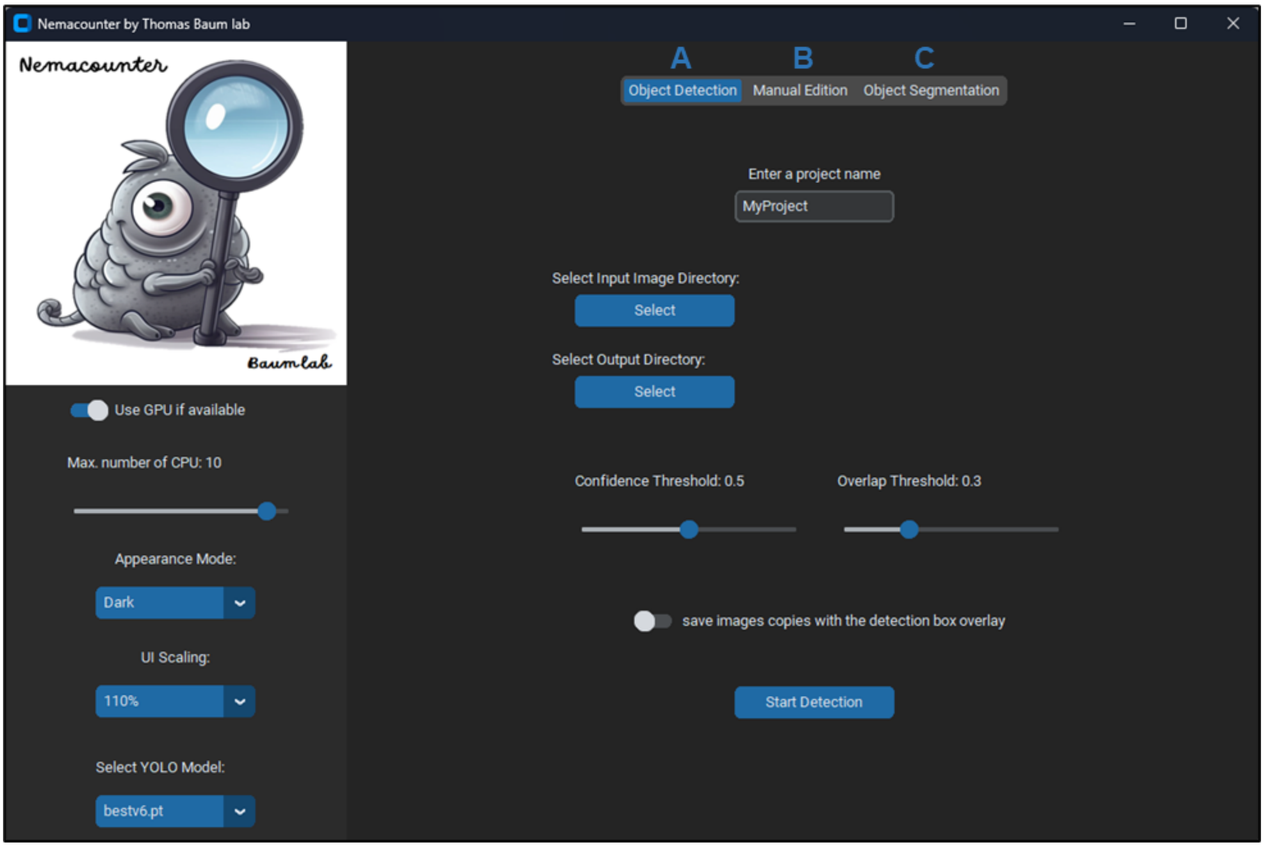

##### Left Sidebar Settings (Applies to All Tabs)

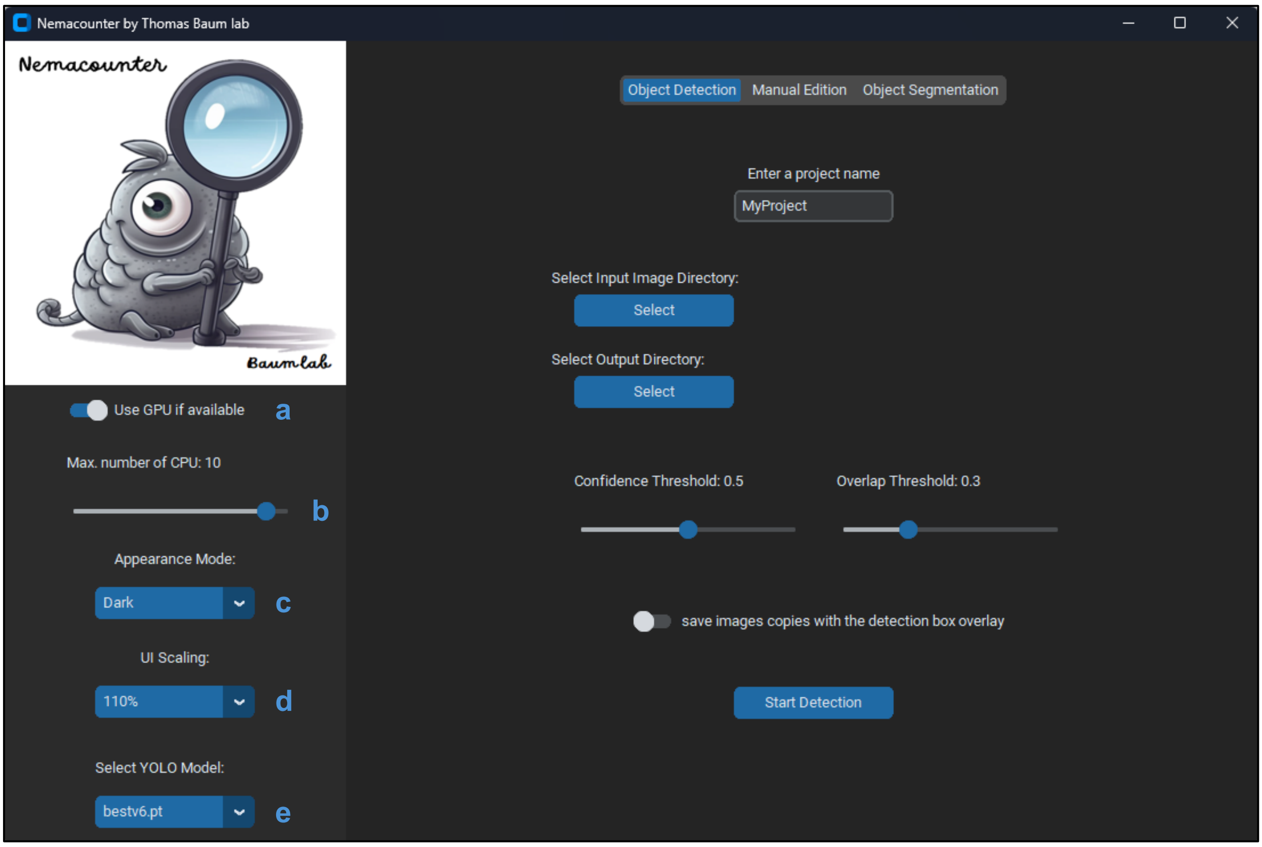


**a. Use GPU if Available**

- **Description**: Toggle the use of GPU for processing if available.
- **How to Use**: Click the switch to enable or disable GPU usage.

**b. Max. Number of CPU**

- **Description**: Set the maximum number of CPU cores for processing.
- **How to Use**: Drag the slider to adjust the number of CPU cores. The current number of selected cores is displayed above the slider.

**c. Appearance Mode**

- **Description**: Change the appearance mode of the software.
- **How to Use**: Click the dropdown menu and select the desired appearance mode (Dark, Light, or System).

**d. UI Scaling**

- **Description**: Adjust the scaling of the user interface.
- **How to Use**: Click the dropdown menu and select the desired scaling percentage.

**e. Select YOLO Model**

- **Description**: Choose the YOLO model (.pt file) for object detection.
- **How to Use**: Click the dropdown menu and select the desired YOLO model file from the list.

**Note**: If a new model.pt is generated by a user, it can be simply added in the "models" folder inside of the "Nemacounter" folder to be selectable in the dropdown menu.

#### Object Detection Tab

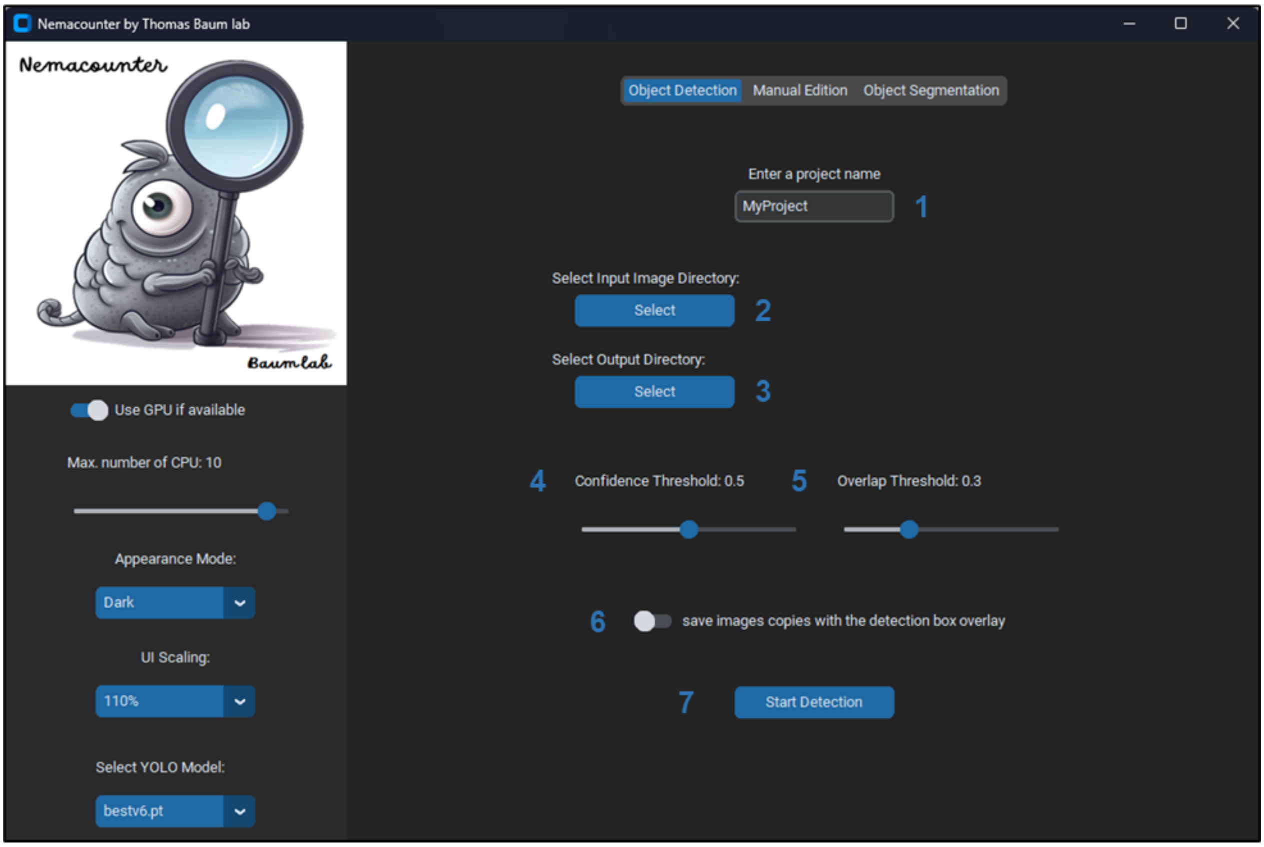


##### 1. Enter a Project Name

- **Description**: Set the name of your project.
- **How to Use**: Click the text box and type the desired project name.

##### 2. Select Input Image Directory

- **Description**: Choose the directory containing the images to be processed.
- **How to Use**: Click the "Select" button and navigate to the desired directory. Click "OK" to confirm.

##### 3. Select Output Directory

- **Description**: Choose the directory where the output results will be saved.
- **How to Use**: Click the "Select" button and navigate to the desired directory. Click "OK" to confirm.

##### 4. Confidence Threshold

- **Description**: Set the confidence threshold for object detection.
- **How to Use**: Drag the slider to adjust the threshold. The current value is displayed above the slider.

5. **Overlap Threshold**

- **Description**: Set the overlap threshold for object detection.
- **How to Use**: Drag the slider to adjust the threshold. The current value is displayed above the slider.

6. **Save Images with Detection Box Overlay**

- **Description**: Enable or disable saving images with detection boxes overlayed.
- **How to Use**: Click the switch to toggle this option on or off.

7. **Start Detection**

- **Description**: Begin the object detection process with the specified settings.
- **How to Use**: Click the "Start Detection" button to start processing.

#### Manual Edition Tab

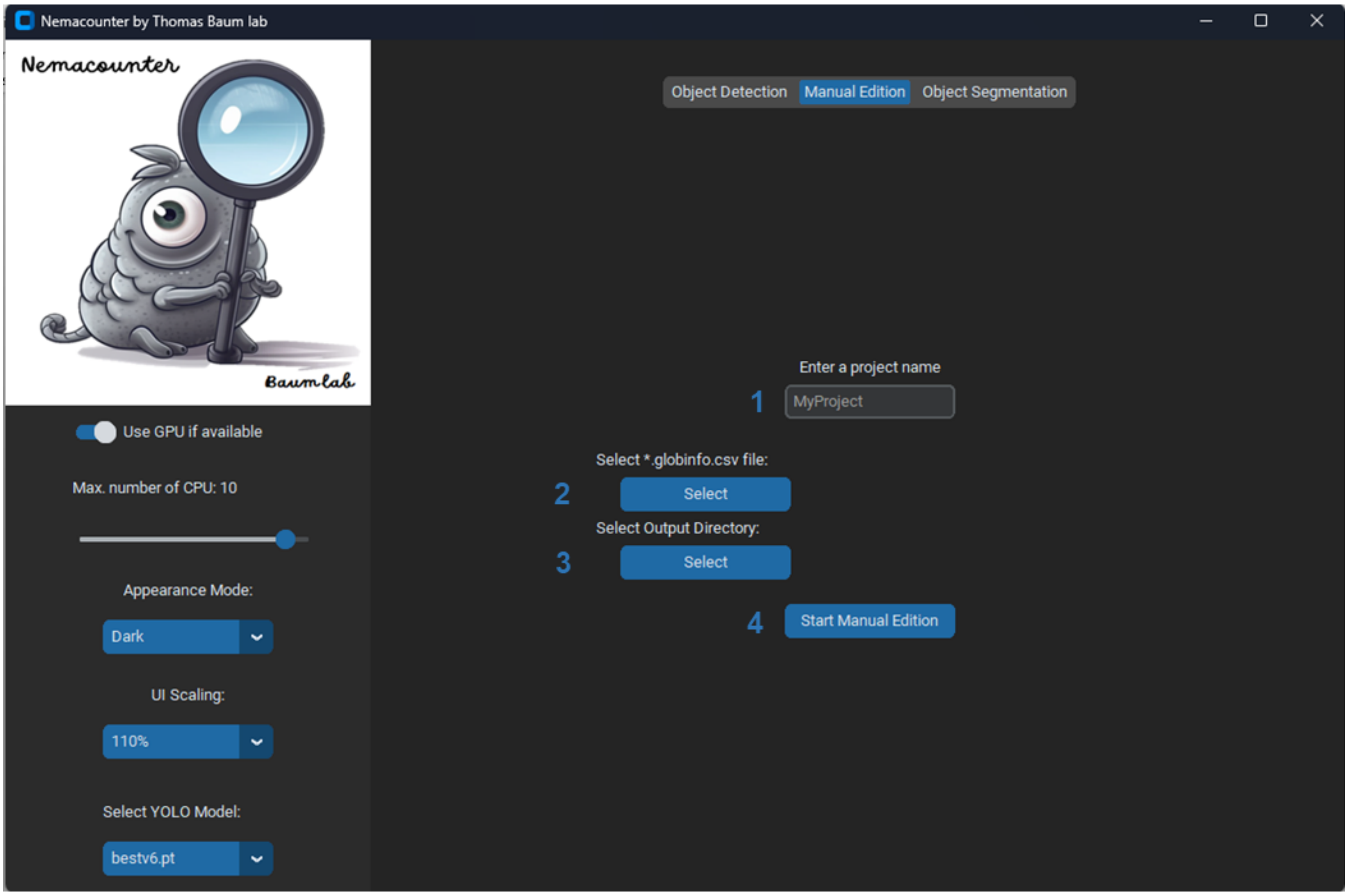


1. **Enter a Project Name**

- **Description**: Set the name of your project.
- **How to Use**: Click the text box and type the desired project name.

2. **Select *.globinfo.csv File**

- **Description**: Choose the CSV file containing the globinfo data generated by the Object Detection tab.
- **How to Use**: Click the "Select" button and navigate to the desired CSV file. Click "OK" to confirm.

3. **Select Output Directory**

- **Description**: Choose the directory where the edited results will be saved.
- **How to Use**: Click the "Select" button and navigate to the desired directory. Click "OK" to confirm.

4. **Start Manual Edition**

- **Description**: Begin the manual edition process with the specified settings.
- **How to Use**: Click the "Start Manual Edition" button to start processing.

#### Object Segmentation Tab

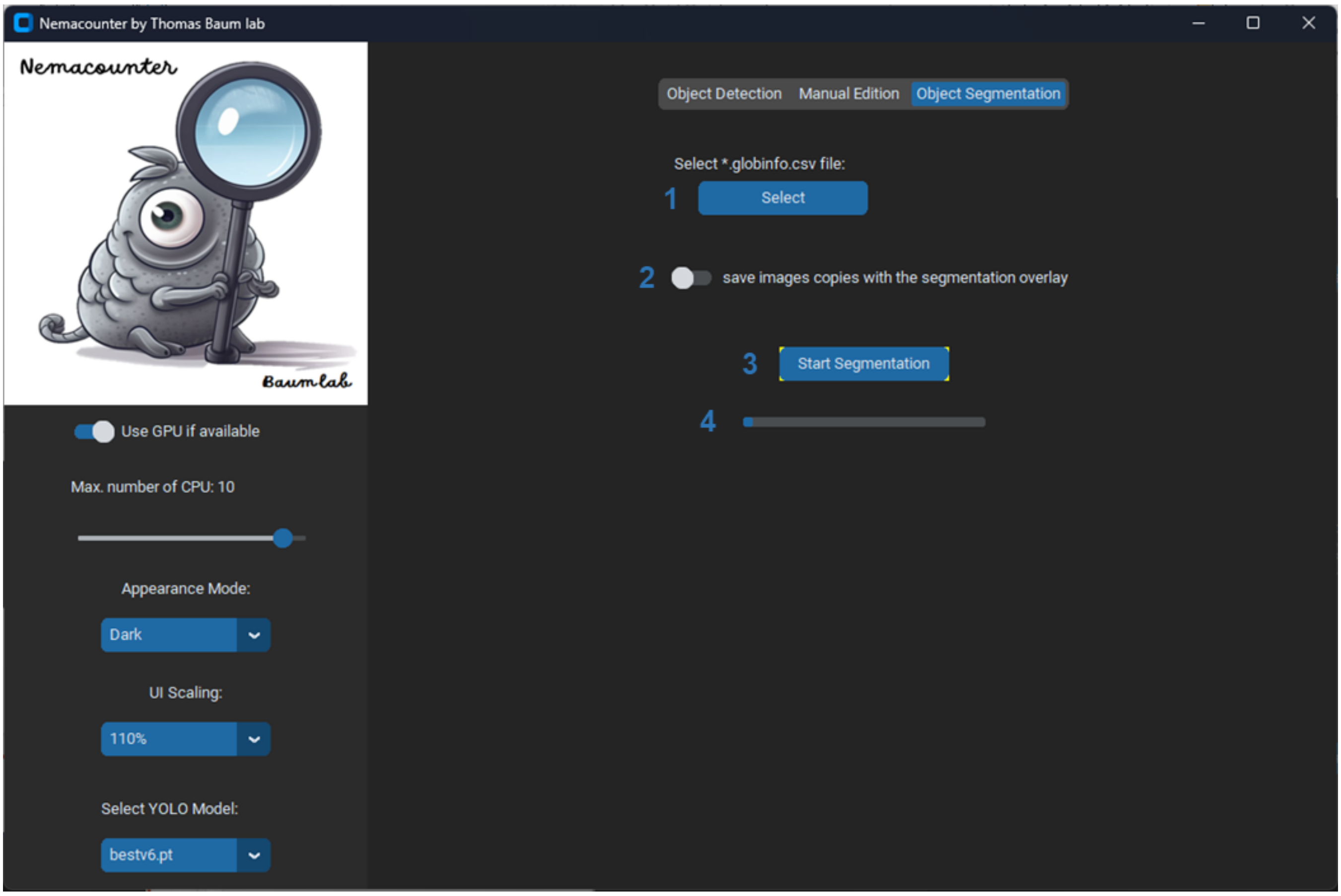


1. **Select *.globinfo.csv File**

- **Description**: Choose the CSV file containing the globinfo data generated by the Object detection tab or the Manual Edition tab.
- **How to Use**: Click the "Select" button and navigate to the desired CSV file. Click "OK" to confirm.

**2. Save Images with Segmentation Overlay**

- **Description**: Enable or disable saving images with segmentation overlays.
- **How to Use**: Click the switch to toggle this option on or off.

**3. Start Segmentation**

- **Description**: Begin the object segmentation process with the specified settings.
- **How to Use**: Click the "Start Segmentation" button to start processing.

**4. Progress Bar**

- **Description**: Displays the progress of the segmentation process.
- **How to Use**: Monitor the progress bar to see the current progress of the segmentation.

---
